# Learning hierarchical sequence representations across human cortex and hippocampus

**DOI:** 10.1101/583856

**Authors:** Simon Henin, Nicholas B. Turk-Browne, Daniel Friedman, Anli Liu, Patricia Dugan, Adeen Flinker, Werner Doyle, Orrin Devinsky, Lucia Melloni

## Abstract

Sensory input arrives in continuous sequences that humans experience as units, e.g., words and events. The brain’s ability to discover extrinsic regularities is called statistical learning. Structure can be represented at multiple levels, including transitional probabilities, ordinal position, and identity of units. To investigate sequence encoding in cortex and hippocampus, we recorded from intracranial electrodes in human subjects as they were exposed to auditory and visual sequences containing temporal regularities. We find neural tracking of regularities within minutes, with characteristic profiles across brain areas. Early processing tracked lower-level features (e.g., syllables) and learned units (e.g., words); while later processing tracked only learned units. Learning rapidly shaped neural representations, with a gradient of complexity from early brain areas encoding transitional probability, to associative regions and hippocampus encoding ordinal position and identity of units. These findings indicate the existence of multiple, parallel computational systems for sequence learning across hierarchically organized cortico-hippocampal circuits.

## INTRODUCTION

We receive continuous input from the world and yet experience it in digestible chunks. In the domain of language, for example, acquisition and use require extracting meaningful sequences such as words, phrases and sentences out of a continuous stream of sounds, often without clear acoustic boundaries or pauses between linguistic elements (*1*). This segmentation ability occurs incidentally and effortlessly and is thought to be a core building block of development. Indeed, young infants can learn transitional probabilities between syllables (*2*) or shapes (*3*) to extract embedded regularities after minimal exposure. In a seminal study (*2*), eight month-old infants segmented words after brief exposure to a continuous sequence of an artificial language in which transitional probabilities between syllables indicated word boundaries. Since this discovery, similar abilities have been demonstrated in adults (*4, 5*), who also rely on transitional probabilities and other statistical properties (*6*-*8*). This behavior—referred to as “statistical learning” (SL)—occurs across many different sensory modalities, tasks, and even species. SL represents a fundamental behavior, and yet the brain mechanisms that support this cognitive function are poorly understood.

Brain regions such as the hippocampus and the inferior frontal gyrus have been implicated in visual (*9, 10*) and auditory SL (*10, 11*). As prior studies have focused on how the brain changes after SL, the role of these brain areas during the *acquisition* of statistical regularities remains largely unexplored. Even less is known about what information is represented in these learned regularities, and whether sequences are encoded similarly or in a complementary fashion across these brain areas. Regularities extracted during SL range from simple to complex, including transitional probabilities between adjacent elements (i.e., uncertainty given a local context), ordinal position in a sequence (i.e., whether an element takes the first, second, third, etc. position), and the identity of the learned unit (i.e., a specific higher-order chunk such as a word) (*12*). Finally, the fact that SL has been observed across sensory modalities raises the question of whether the same brain areas and algorithms support extraction and representation of regularities (*13*).

To answer these questions, we collected intracranial recordings (ECoG) from 23 human epilepsy patients with broad cortical and hippocampal coverage during an SL task. We used neural frequency tagging (NFT) (*14, 15*) to identify recording sites responsive to the underlying regularities of the SL stimuli over different timescales (e.g., syllables and words). Combining ECoG and NFT, we describe the location and temporal tuning of the neural response. Following identification of responsive sites, we used representational similarity analysis (RSA) to determine which aspect(s) of the temporal regularities are represented i.e., transitional probabilities, ordinal position and identity. Finally, we related the neural circuits, online dynamics and representational changes for SL across auditory and visual modalities.

We found that SL occurs quickly in both auditory and visual modalities. In both modalities, partially overlapping neural circuits encoded statistical units (e.g., words, fractal pairs) and their constituent sensory elements (e.g., syllables, images). This learning was supported by rapid changes in the similarity space of neural representations, with structure encoded at multiple levels: (*1*) transitional probabilities, with elements grouped by probability strength; (*2*) ordinal position, with elements grouped by sequence order; and (*3*) identity, with elements grouped by unit in which they are embedded. Auditory and visual elements underwent similar SL-related representational changes, yet involved brain areas only partially overlapped (generally supramodal areas, such as inferior frontal gyrus, anterior temporal lobe, and the hippocampus). These results provide mechanistic insight into a fundamental human learning ability, revealing how cortical areas respond to the structure of the world. Our findings also highlight NFT as a versatile tool for investigating incidental learning in preverbal infants and other non-verbal patient populations.

## RESULTS

### Behavioral evidence of auditory statistical learning

To investigate the neural circuits and computations underlying SL, we presented a group of 17 epilepsy patients with brief (2 min x 5 blocks) auditory streams of syllables in which the structure of the sequence was manipulated. In structured streams, each syllable was placed into the first, second, or third position of a three-syllable word or “triplet” (**Fig. 1A**). A continuous stream of syllables was generated by randomly inserting multiple repetitions of each word without pauses or prosodic cues between words. In random streams, syllables were inserted the same number of times but in a random order at the syllable level. Thus, the transitional probabilities were low and uniform, without a word-level of segmentation.

**Fig. 1.**
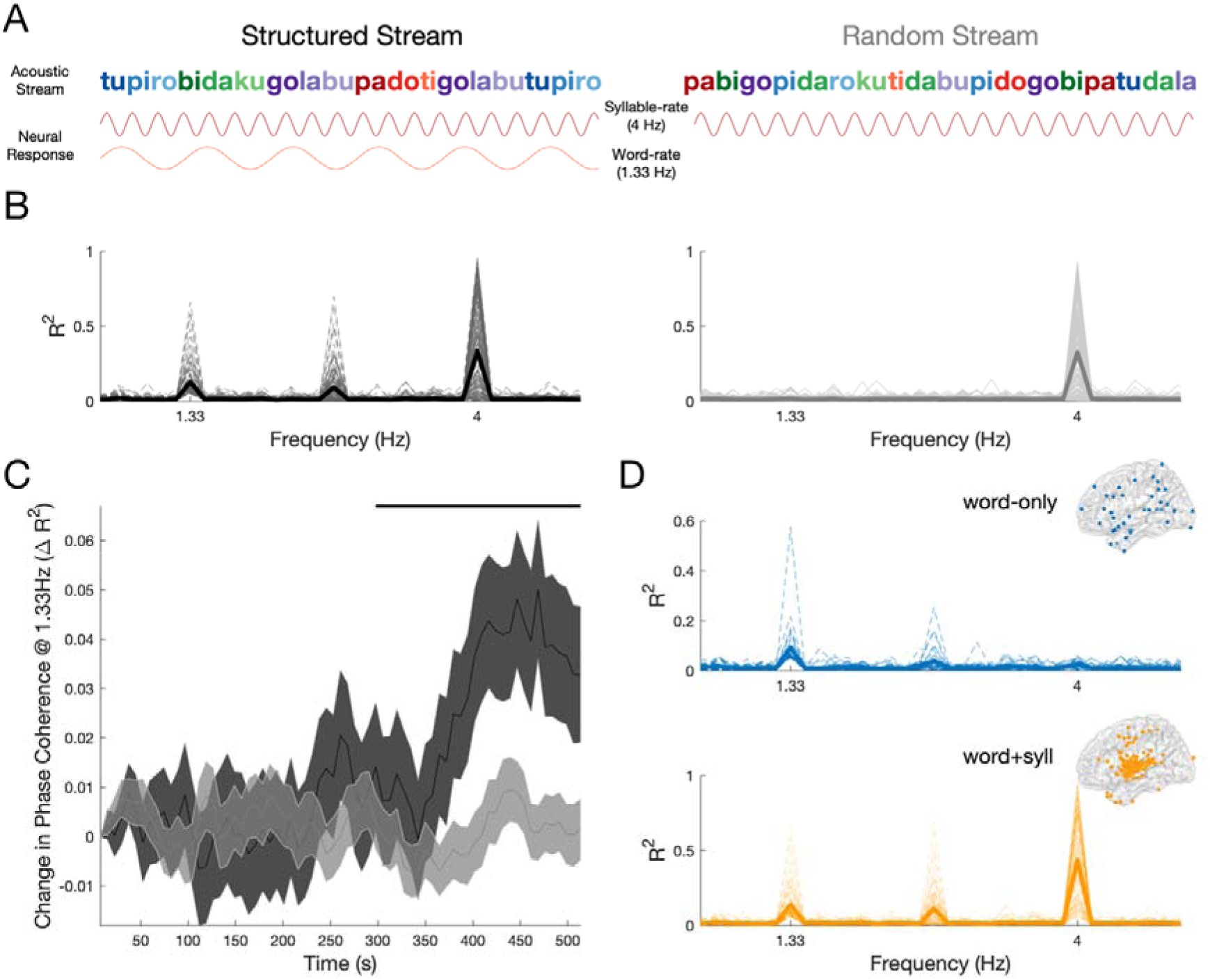
Neural tracking of auditory statistical learning. **(A)** Schematic depiction of the auditory SL task. The structured stream (left) contained 12 syllables (250 ms SOA, 4 Hz) in which the transitional probabilities formed four words (color coded for visualization, 750 ms SOA, 1.33 Hz). The random stream (right) contained the same 12 syllables in a random order. The predicted neural response is shown below each syllable stream: syllable tracking (top) was expected in both conditions, whereas word tracking (bottom) was expected only in the structured condition. **(B)** Phase coherence spectrum in neural data for the structured (left, black) and random (right, gray) conditions from 1,898 electrodes in 17 patients. Each electrode is depicted with a thin line and the average with a thick line. **(C)** Timecourse of coherence response for all electrodes showing word-rate tracking (1.33Hz) for the structured (black) and random (gray) conditions. Solid line above the timecourse indicates when the change in coherence increased above zero (p<0.05, one-sided cluster-corrected permutation test). (**D)** Phase coherence spectrum in the structured condition for electrodes showing word-tracking responses, in two groups: electrodes that tracked words only (top, blue) and electrodes that tracked both words and syllables (bottom, orange). Inset shows the localization of word-only (blue) and word+syll (orange) electrodes.

Participants were blinded to the stimulus structure. They were asked to perform a 1-back cover task, in which they had to detect occasional repetitions of individual syllables that had been inserted into both stream types (*16*). This task has been used to evaluate SL online while assuring attention to the SL stimulus. Accuracy in both streams was high and not statistically different (t(*16*)=2.03, p=0.06), indicating that participants attended to the stimuli across both the structured (mean d’ = 1.04, t(*16*) =14.41, p<0.01) and random (mean d’=0.87, t(*16*)=14.27, p<0.01) streams. Critically, we found that behaviorally, reaction times to repeated syllables in the structured stream (mean=733 ms) were significantly faster than in the random stream (mean=917 ms; Z=-3.3, p=0.001, fig. S1), suggesting that facilitation had occurred due to learning of the underlying structure.

Immediately after exposure to both streams, participants were informed of the hidden structure and were asked to perform an explicit recognition task. Recognition of the hidden words in the structured stream was assessed using a 2-alternative forced choice (2AFC) task between the hidden words and part-words. Part-words consisted of previously shown sequences of syllables but that spanned words and thus had overall lower transitional probabilities. Offline explicit recognition of the hidden words did exceed chance performance in some subjects (50%; mean=45.4%, SD=10%; Z=-1.84, p=0.07).

The same procedure was used in a separate cohort of healthy subjects in which we replicated the online incidental learning effect in the reaction times i.e., faster responses to syllable repetition in the structured than in then random condition (mean structured = 625 ms, mean random = 828 ms, Z=-3.8, p<0.001, N=18). Offline explicit recognition was significantly better than chance in this neurotypical cohort (50%; mean=57.3%, Z=2.04, p=0.04, **fig. S1**).

### Neural tracking of auditory statistical learning

We obtained direct neurophysiological signals from 1,898 intracranial electrodes in the 17 participants, comprehensively covering the frontal, parietal, occipital, temporal lobes and the hippocampus in both hemispheres (**fig. S2**). We capitalized on NFT to evaluate the temporal dynamics of the neural activity in order to scout for cortical areas responding at the rate of the learned regularities. The sensitivity of NFT to track SL has been previously demonstrated using non-invasive techniques, i.e., EEG and MEG (*17, 18*) enabling us to definitively resolve the cortical areas exhibiting selective temporal tuning to the learned regularities. Specifically, NFT was used to track representations of segmented units at two hierarchical levels of the stream (*14, 15, 17*). Entrainment at the syllabic frequency (4 Hz) should be present in both structured and random streams; while entrainment at the word-level frequency (corresponding to triplet boundaries or 1.33 Hz) should emerge during exposure to the structured but not random stream.

We first evaluated within-electrode phase coherence in the field potential (FP)(*15*) for the structured and random streams, respectively. Consistent with our hypothesis, there was a significant peak in the phase coherence spectrum at the syllable rate (i.e., 4 Hz) for both structured and random streams (p<0.05, FDR corrected). In addition, there was a significant peak at the word rate (i.e., 1.33 Hz), but only for the structured stream (p<0.05, FDR corrected; **Fig. 1B**). There was also a significant phase coherence peak at 2.66 Hz for the structured stream (p<0.05, FDR corrected). This may reflect an oscillation at the rate of syllable *pairs*, consistent with evidence that participants can learn sequential pairs embedded in triplets, in addition to the triplets themselves (*19*); alternatively, this may be a harmonic of the word rate. The word-rate response in the structured stream emerged rapidly within 260 seconds (exceedance mass: sum(*T*) = 98, p= 0.003) and increased over time (**Fig. 1C**); while no increase in the strength of the word-rate response was observed for the random stream, ruling out effects of endogenous entrainment over time unrelated to learning. Coherence at the word rate in the structured stream was replicated across participants with 16/17 patients exhibiting significant entrainment in at least one electrode; by contrast, no electrodes showed entrainment at the word rate for the random stream. This finding further supports NFT as a sensitive and robust tool for assessing online SL.

We then exploited the unique spatial resolution afforded by ECoG to localize which cortical areas became synchronized to the word rate in the structured condition (**Fig. 1D**). Different temporal tuning responses were observed across electrodes. One tuning profile corresponds to electrodes that tracked both words and syllables (word+syll). These were located primarily in the superior temporal gyrus (STG), with smaller clusters in motor cortex and pars opercularis. The other tuning profile reflected electrodes which exclusively tracked the words (word-only). These were located in inferior frontal gyrus (IFG) and anterior temporal lobe (ATL) (**table S1**). These functional responses indicate temporal selectivity to both the input (in this case the syllable) and higher-order learned units (for word+syll electrodes), or to higher-order learned units alone without responding to the acoustic features of the input conveying the structure (for word-only electrodes). This organization reflects the neuroanatomy of the auditory processing hierarchy, with lower-order function in STG and higher-order function in surrounding fronto- and temporo-parietal cortex (*20*). Thus, we reasoned that word-only responses may arise from higher-level stages of processing than word+syll responses. To quantify this anatomical grouping by electrode type, we tested the hypothesis that electrodes belonging to one type (i.e., word-only or word+syll) tend to group together (e.g., nearest electrode was of the same type) using a Bayesian binomial test. Bayesian analysis provided evidence in support of this hypothesis (nearest electrode in ‘same-type’ vs. ‘different type’, log(BF10)=40.75).

We also conducted the same analysis with the high-gamma band (HGB). The HGB responses is thought to reflect multiunit firing (*21*) and considered more selective and spatially confined compared to the FP (*22*). We observed the same two types of responses and the topographies for the two types resemble those described above for FP (**fig. S3**).

### Representational analysis in auditory statistical learning

The results so far provide evidence of segmentation of the continuous auditory stream with characteristic tuning in lower-order areas in STG and higher-order areas in surrounding fronto- and temporo-parietal cortex. But what is driving this segmentation? The neural response to segmentation could be based on at least three statistical cues in the stream: *transitional probabilities* (within word, 1.0; between word, 0.33), *ordinal position* (1^st^, 2^nd^, or 3^rd^ position), or *word identity* (blue, green, purple, or red word, as in colors from **Fig. 1A**). Although all three cues could be used to mark the start and end of the words and thus drive segmentation, they differ in content and facilitate unique cognitive functions. For instance, coding based on transitional probabilities and the entailed difference in entropy between high and low TP can serve as a strong prediction error cue to drive attention and segmentation. Coding of ordinal position represents a flexible and abstract code allowing the recombination of elements and might explain previous findings on phantom words (*23*), whereby subjects accept as legal, strings that have never appeared during the exposure phase as long as ordinal position is preserved. Yet, only coding based on identity gives access to individual words which can then be mapped onto meaning (*24*).

To evaluate what information is being represented we used a multivariate pattern similarity approach. In the case of word identity, for example, we reasoned that SL would change the representational space of stimuli such that syllables belonging to the same word would evoke more similar neural activity patterns across electrodes (*9*); this clustering could in turn provide a basis for segmentation (*7*). Alternatively, the neural representations of syllables may cluster by ordinal position or transitional probability, allowing us to test which of these cues was learned and whether similar or complementary codes are observed across brain areas. We quantified the representational space of syllables from FP separately within the sets of electrodes identified as exhibiting word-only and word+syll coherence. We focused on electrodes exhibiting such tuning in the HGB, given previous reports of its correlation to neural spiking (*21*). In addition, we separately investigated neural representations across electrodes in the hippocampus, as previous studies have shown that the hippocampus is necessary for robust SL (*10, 25*). We calculated the correlation distance between the patterns of neural activity across electrodes within each set of electrodes (word-only, word+syll, and hippocampus), for each pair of syllables, and applied multidimensional scaling (MDS) to visualize the similarity structure.

We found that the three sets of electrodes encoded different information: For word+syll electrodes, MDS of the distances between syllables revealed a representation of transitional probabilities **(Fig. 2A)**, grouping syllables based on whether their probability given the preceding syllable was low (1^st^) vs. high (2^nd^, 3^rd^). In contrast, word-only electrodes represented ordinal position **(Fig. 2B)**, grouping syllables based on which position they occupied in the words (1^st^ vs. 2^nd^ vs. 3^rd^), as well as based on the identity of the word to which they belong (**Fig. 2B**, dimension 2). Finally, hippocampal electrodes showed grouping by word identity only **(Fig. 2C)**. The clustering of responses by transitional probability, ordinal position and identity is consistent with fast learning during exposure as they were absent during the first block (∼2 mins) and present by the fifth block **(fig. S4)**. Moreover, no clustering of syllable representations was observed in any electrode set when the same analysis was performed on the random stream (**fig. S5**). This demonstrates that changes in representational space for the structured stream resulted from SL and not properties of the individual stimuli per se.

**Fig. 2.**
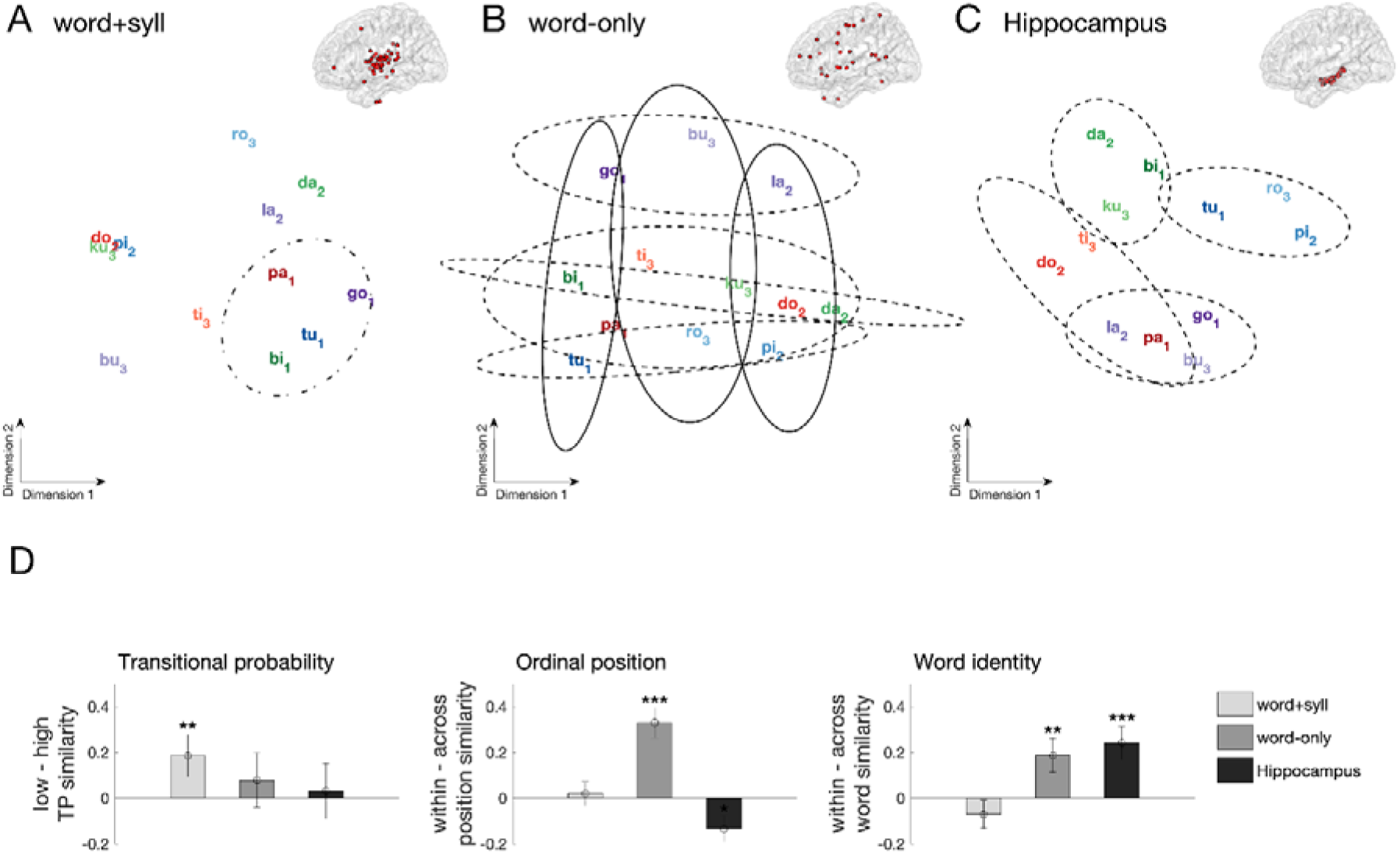
Pattern similarity results during auditory statistical learning. Multidimensional scaling (MDS) distances between syllabic responses across electrodes showing significant **(A)** word+syll responses and **(B)** word-only responses, as well as **(C)** across electrodes from the hippocampus. Individual words are color-coded, subscripts represent ordinal position (e.g., “tu_1_pi_2_ro_3_”). Dot-dashed ellipses indicate grouping by transitional probability, solid ellipses outline grouping by ordinal position, and dashed ellipses indicate grouping at the level of the individual words (color coded). **(D)** Comparison of multivariate similarity for syllables in the auditory SL task. **Left:** similarity by transitional probability. Greater within-class similarity indicates stronger grouping of syllables with low transitional probability (0.33) than syllables with high transitional probability (1.0). A Friedman test indicated a main effect of electrode type on TP similarity (χ^2^ = 22.03, p<0.001). **Middle:** within vs. between similarity for ordinal position. Greater within-class similarity indicates stronger grouping of syllables holding the same 1^st^, 2^nd^, or 3^rd^ position in a word. A Friedman test indicated a significant main effect of electrode type (χ^2^ = 790.35, p<0.001). **Right:** within vs. between similarity for word identity. Greater within-class similarity indicates grouping of syllables into individual words. A Friedman test indicated a significant main effect of electrode type (χ^2^ = 265.29, p<0.001). (***p<0.001, **p<0.01, Wilcoxon rank sum test, error bars denote the population standard error of the mean).

To evaluate statistically what information is encoded in each set of electrodes, we compared pattern similarity across syllables for grouping consistent with transitional probability, ordinal position and unit identity. To that end, we compared pattern similarity for same versus different classes i.e., similarity of syllables with low vs. high transitional probability, same vs. different ordinal position, and same vs. different word identity, for each electrode types (word-only, word+syll, hippocampus). Consistent with the MDS, low transitional probability coding was only observed for word+syll electrodes (**Fig. 2D, left**). Reliable coding for ordinal position was observed for the word-only electrodes **(Fig. 2D, middle)**. Word identity was observed both in word-only electrodes and in the hippocampus **(Fig. 2D, right)**. These results show that even brief exposure to auditory regularities can reshape the representational space of syllables throughout cortex and the hippocampus, giving rise to clustered neural representations along several dimensions. That is, learning of sequences shapes representations at multiple levels concurrently, with a division of labor across lower- and higher-order brain areas in terms of simple and generic vs. complex and specific regularities.

### Behavioral evidence of visual statistical learning

Segmenting continuous input into discrete units extends also to stimuli in the visual domain, e.g., to build representations of scenes and events (*26*). Controversy remains as to whether similar SL mechanisms are engaged in segmenting and acquiring structure across auditory and visual domains (*13*). To investigate whether similar coding principles could be at work in the visual modality, we tested visual SL in 12 intracranial patients. These participants were exposed to brief (2 min x 5 blocks) visual streams of fractal images (375 ms each) in which the structure of the sequence was manipulated (*9*). In the structured streams, each fractal was assigned to the first or second position of a pair (**Fig. 3A**). We generated a continuous stream of fractal pairs by randomly inserting each pair without breaks or other cues between pairs. In the random streams, fractals were inserted the same number of times but in a random order at the fractal level. As a result, there were no pairs to segment.

**Figure 3.**
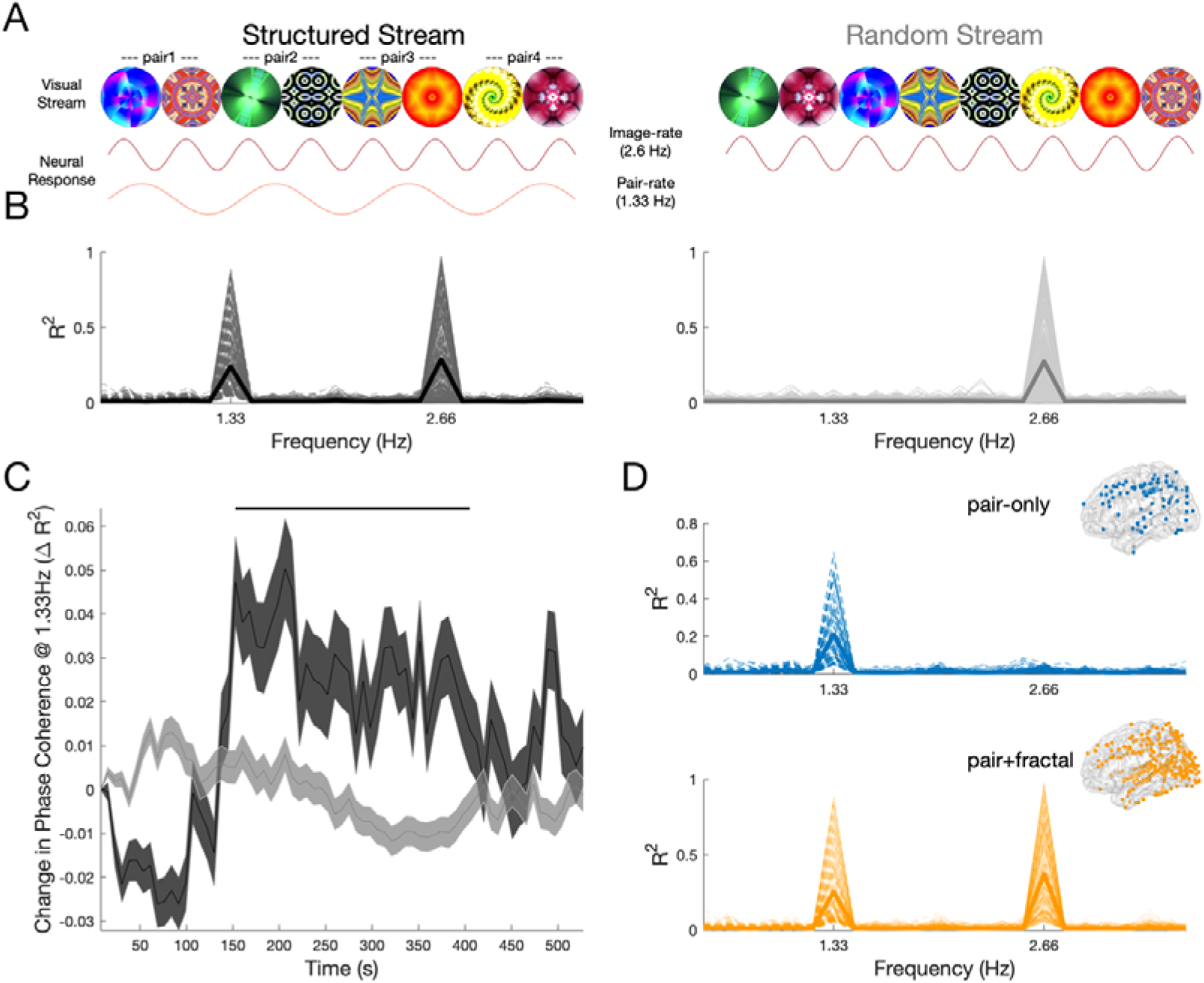
Neural tracking of visual statistical learning. **(A)** Schematic depiction of the visual SL task. The structured stream (left) consisted of a continuous visual stream of eight fractals (375 ms SOA, 2.66 Hz). The transitional probabilities were adjusted to form four fractal pairs (750 ms SOA, 1.33 Hz). Note that the SOA of the fractals was elongated compared to the syllables to match the frequency of the learned units (pairs and words), given that there were two fractals per unit and three syllables. The random stream (right) contained the same fractals but in random order. The predicted neural responses are shown under each stream: fractal tracking is expected for both streams while pair tracking is expected for the structured stream only. **(B)** Phase coherence spectrum in neural data for the structured (left, black) and random (right, gray) conditions from 1606 electrodes in 12 patients. Each electrode is depicted with a thin line and the average with a thick line. **(C)** Timecourse of coherence response for all electrodes showing pair-rate tracking (1.33Hz) in the structured (black) and random (gray) conditions. Solid line above the timecourse indicates when the change in coherence increased above zero (p<0.05, one-sided cluster-corrected permutation test). **(D)** Phase coherence spectrum in the structured condition for electrodes showing pair-tracking responses, in two sets: electrodes that tracked pairs only (left, blue) and electrodes that tracked pairs and fractals (right, orange). Inset shows the localization of pair-only (top, blue) and pair+fractal (bottom, orange) electrodes.

As in the auditory sequence, participants were not informed about the presence of structure in some of the sequences. Instead, they were asked to perform a 1-back cover task, in which they had to detect repetitions of individual fractals that had been occasionally inserted into both stream types (*16*). Accuracy was similarly high in both structured (mean d’= 1.66, *t*(*11*) =5.27, p<0.001) and random (mean d’=1.78, *t*(*11*)=6.96, p<0.001) streams, and did not differ (*t*(*11*)=-0.46, p=0.65). This suggests that participants were equally engaged and attentive during both streams. Consistent with incidental SL of the structure, we again found significantly faster reaction times in the structured stream (mean=621 ms) than in the random stream (mean=729 ms; Z =-1.99, p=0.04, fig. S6).

Following exposure to both streams, participants performed a 2AFC recognition task to assess explicit learning of the fractal pairs. Offline explicit recognition was at chance performance in these participants (50%; mean=53.3%, SD=8%; *Z* = 1.23, p=0.22). However, as with auditory SL, we again replicated the findings in a neurotypical sample and found evidence of incidental leaning in the reaction times (e.g. faster reaction times in the structured condition; mean structured = 630 ms, mean random = 648 ms; Z=-2.1, p=0.03, N=14), while offline, explicit recognition was significantly better than chance (50%; mean=57.7%, SD=10%; Z=2.45, p=0.01, N=14, fig. S6) in this cohort.

### Neural tracking of visual statistical learning

We next turned to NFT to identify brain areas exhibiting SL in neurophysiological recordings from 1606 intracranial electrodes in the 12 patients, extensively covering frontal, parietal, temporal and occipital cortex (**fig. S2**). Specifically, we expected an entrainment response at a 2.66 Hz frequency to individual fractals and at a 1.33 Hz frequency to the learned pairs, the latter only for the structured stream. Providing evidence for the acquisition of regularities, we observed a significant peak in the phase coherence spectrum at the pair rate (i.e., 1.33 Hz) but only for the structured stream (p<0.05, FDR corrected; **Fig. 3B**). A significant peak at the fractal rate (i.e., 2.66 Hz) was found for both structured and random streams (p<0.05, FDR corrected).

Coherence at the pair rate was replicated across subjects with 12/12 patients exhibiting entrainment in at least one electrode. Increases in pair-rate responses were observed within 160 s of exposure (exceedance mass: sum(*T*) = 103, p= 0.004), subsequently plateauing despite further exposure to the structured stream. Such an increase in the pair-rate response was absent in the random stream ruling out spurious entrainment as a function of time (**Fig. 3C**).

As in the auditory SL, we observed an anatomical and hierarchical segregation between two temporal tuning profiles of electrodes: one showing entrainment at the fractal and pair rates (pair+fractal, **Fig. 3D, bottom**) and clustered mostly within occipital (striate and extrastriate) and parietal cortex (IPS); the other showing entrainment at the pair-rate only (pair-only, **Fig. 3D, top**) localized more anteriorly in frontal (middle and superior), parietal and temporal cortex (**table S1**). The same separation between pair+fractal and pair-only responses and spatial arrangement across electrodes was observed when restricting the analysis to the HGB (**fig. S3**).

### Representational analysis in visual statistical learning

In the auditory modality we observed representational changes indicative of sequence learning based on transitional probabilities, ordinal position and word identity across sets of electrodes. How do visual regularities shape neural representations? As learned units in the visual modality contained only two elements (i.e., pairs), grouping based on transitional probabilities and ordinal position yield similar results — both cues predict grouping of the first fractal in each pair with the first fractals of other pairs and grouping of the second fractal with the other second fractals. However, although both transitional probability and ordinal position depend on grouping of first fractals together and second fractals together, the representational impact of these cues can be quantified in different, non-exclusive ways (see below). Thus, in the following, we refer to this grouping as consistent with either transitional probability or ordinal position, in contrast with pair identity, which predicts that the first and second members of each pair will be grouped together and different from the other pairs (*9*). Follow-up quantification will allow to differentiate grouping based on transitional probability, ordinal position and identity.

We conducted multivariate pattern analysis on the FP separately for the sets of electrodes showing pair+fractal responses, pair-only responses, in addition to electrodes in the hippocampus. For the pair+fractal and pair-only responses we selected electrodes demonstrating NFT in the HGB. We calculated the correlation distance between the spatial patterns of neural activity across electrodes for every pair of fractals for each of the three electrode sets. We again found that the three sets of electrodes encoded different information. For the pair+fractal electrodes, multidimensional scaling of the distances between fractals revealed a representation consistent with transitional probabilities and/or ordinal position (representation of first vs. second, **Fig. 4A**). Replicating the auditory findings, for pair-only electrodes we found concurrent grouping for transitional probabilities and/or ordinal position (first dimension) and grouping for pair identity (second dimension) (**Fig. 4B**). In the hippocampus, however, grouping was only by pair identity **(Fig. 4C)**. Clustering was not observed in any electrode set for the random stream (**fig. S7**) and consistent with fast learning during exposure the clustering emerged by the fifth block but was absent during the first block (**fig. S8**). These results indicate rapid changes in representational space as a function of visual SL.

**Fig. 4.**
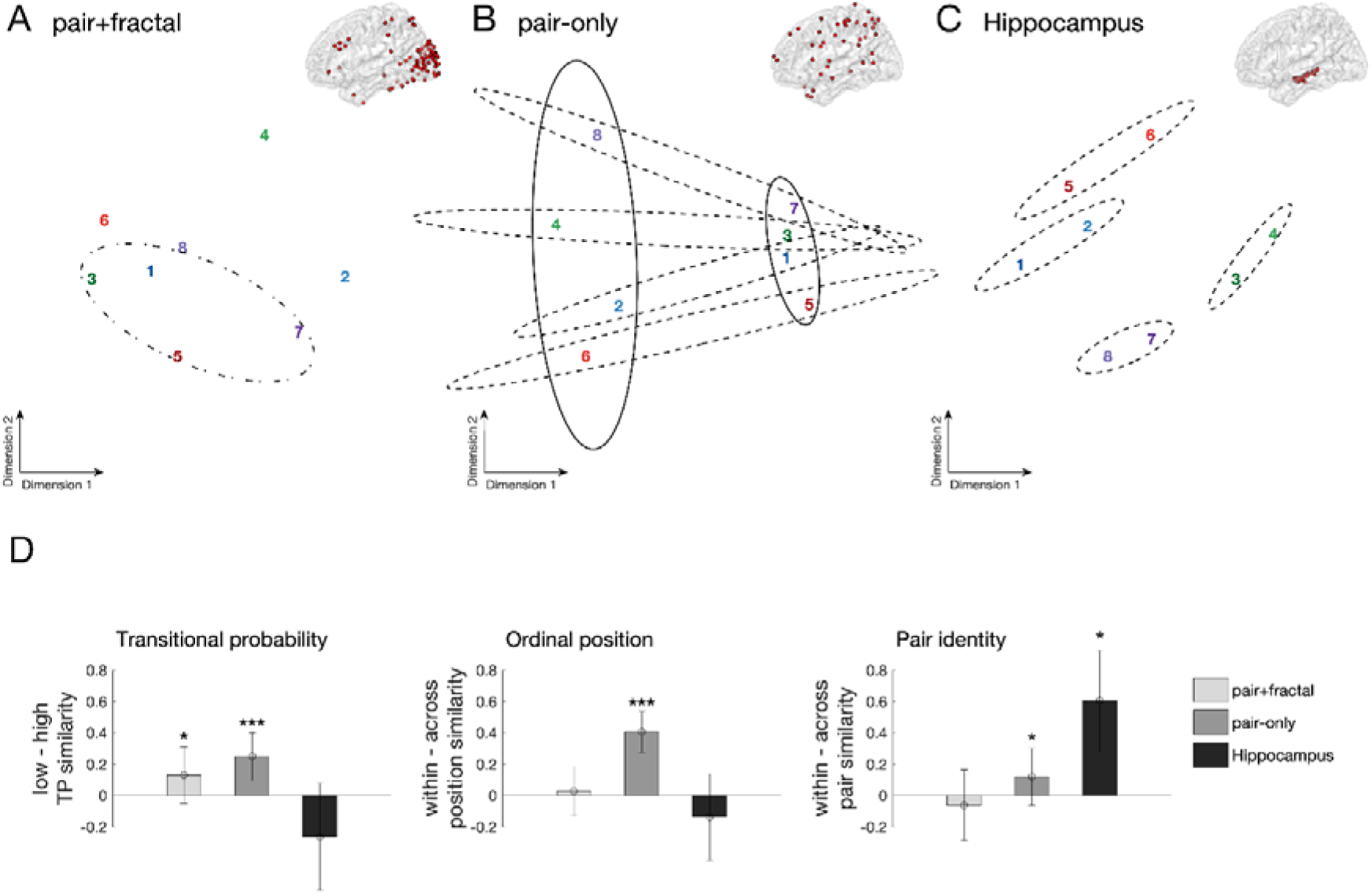
Pattern similarity results during visual statistical learning. Multidimensional scaling (MDS) of distances between responses to individual fractals across (**A**) pair-only, (**B**) pair+fractal, and (**C**) hippocampal electrodes. Pairs are color-coded, odd numbers refer to the first position, even numbers to the second position. Dot-dashed ellipses outline grouping by transitional probability/ordinal position in pair+fractal electrodes. Solid ellipses outline grouping by transitional probability/ordinal position in pair-only electrodes. Dashed ellipses indicate grouping by pair in pair-only and hippocampal electrodes. (**D**) Comparison of multivariate pattern similarity for fractals in the visual SL task. **Left:** within vs. between similarity for low vs. high transitional probability. Greater within-class similarity indicates stronger grouping of fractals with a low transitional probability (0.33) over fractals with a high transitional probability (1.0). A Friedman test indicated a main effect of electrode type on TP similarity (χ^2^ = 19.3, p<0.001). **Middle:** within vs. between similarity for ordinal position. Greater within-class similarity indicates grouping of fractals holding the same 1^st^ or 2^nd^ position in a pair. A Friedman test indicated a main effect of electrode type (χ^2^ = 122.2, p<0.001). **Right:** within vs. between similarity for pair identity. Greater within-class similarity indicates grouping of fractals into pairs. A Friedman test indicated a main effect of electrode type (χ^2^ = 40.04, p<0.001). (***p<001, *p<0.05, Wilcoxon rank sum test, error bars denote the population standard error of the mean).

To statistically evaluate the groupings, we collapsed pattern similarity across fractals belonging to different classes for each electrode set (pair+fractal, pair-only, and hippocampus). This allowed us to quantify the representational impact of each coding scheme. Transitional probability was examined by comparing pattern similarity among first fractals (first-first) with low transitional probability (i.e., relatively unpredictable given preceding fractal) vs. among second fractals (second-second) with high transition probability (i.e., predictable given preceding fractal). Ordinal position was examined by comparing pattern similarity within the same position (first-first, second-second) vs. between different positions (first-second). Pair identity was examined by comparing pattern similarity within pair (e.g., first_1_-second_1_) vs. between pair (first_1_-second_2_).

In line with the MDS, we observed complementary coding across the three sets of electrodes. Pair+fractal electrodes showed greater similarity for low transitional probability but not for ordinal position or identity. In contrast, pair-only electrodes showed greater similarity for fractals with low transitional probability (**Fig. 4D, left**), reliable coding for ordinal position (**Fig. 4D, middle**), and also reliable coding for pair identity (**Fig. 4D, right**). Finally, hippocampal electrodes exclusively showed coding for pair identity.

## DISCUSSION

Using intracranial recordings in humans, we have described how the brain tracks and learns structure within sensory information. SL is accompanied by rapid changes in neural representations, reflected in two functionally and anatomically distinct responses: brain regions tracking lower-level sensory input (i.e. syllables and fractals) and higher-order units (i.e. words and pairs), and brain regions only representing learned higher-order units (i.e. words and pairs). These distinct responses reveal a hierarchical arrangement: the former maps onto early, sensory processing stages (e.g., STS and occipital cortex), while the latter encompasses late, amodal processing stages (e.g., IFG and ATL). In other words, while early processing is domain-specific, late processing is domain-general. Remarkably, these nested structures within sensory streams are extracted and represented in the brain in as little as ∼2 min, consistent with previous behavioral studies (*2*), and even when subjects are not aware of the process.

Our results are consistent with prior work demonstrating how the cortical hierarchy integrates information over increasingly longer temporal windows (*22, 27*). Yet, they go beyond topographical mapping of temporal receptive fields. First, we show how SL shapes the neural representational space within these areas profoundly and rapidly. This contrasts with the much more gradual representational change that occurs over development or with longer-term perceptual learning. Second, we discovered that qualitatively different aspects of sequence knowledge are encoded across different brain areas: sites representing the sensory input *and* higher-order units encode local and generic aspects of sequences, such as their transitional probabilities (or degree of uncertainty). In contrast, sites exclusively representing higher-order structure encode global and also more specific aspects of the sequences such as the ordinal position of the elements, but most importantly the specific identity of the learned unit.

Prior studies on SL suggest that an increase in predictive uncertainty serves as the primary cue for event segmentation. Our results extend this body of work demonstrating that SL also involves the acquisition of higher-order sequence knowledge, i.e., ordinal position and identity. Higher-order structure or “chunks” may serve as the mental units for mapping segmented word forms onto novel word referents (*24*). These results are in line with the hypothesis that the output of the word boundary discovery may provide cues to edges of constituents which in turn can serve as scaffolding for the subsequent discovery of internal structure, i.e., which elements are contained and in which positions (*28*).

Our finding that sequences are represented at multiple levels, from simple and generic to complex and specific regularities, may reconcile two opposing theoretical models in SL. The ‘statistical model’ posits that learners represent statistical relations between elements in the input and do not explicitly represent statistically coherent units in memory. In contrast, ‘chunking models’ posit that learners represent statistically coherent units of information from the input in memory, such that the stored representations are discrete chunks of information. So far, these models have been only contrasted at the behavioral level, e.g., by studying sensitivities to illusory or embedded units (*29, 30*). An intriguing possibility is that these two models actually coexist and map onto the different networks and sequence representations that we report here — i.e., simple representations encoding transitional probabilities and complex representations encoding positional information and unit identity. Previously reported discrepancies in behavioral results may reflect the differential engagement of these two neural processes across different tasks. Alternative mechanisms for SL can also be tested using the RSA approach. In particular, two novel theories can be tested. One posits that SL reflects changes in the similarity space and that transitions are then learned as trajectories through that space (*31*). Another account conceives SL as acquiring a community structure in a symmetric graph with uniform transitional probabilities which are captured by changes in representational similarity (*7*).

The main organizational principles of neural changes underlying SL are shared across the auditory and visual domain. We observed a similar functional clustering of responses, i.e., sensory input + higher-order units and higher-order units-only, in both the auditory and the visual modalities (compare **Fig. 1** with **Fig. 3**). In addition, no clear hemispheric lateralization was observed for the auditory or visual SL in any of the electrodes following higher-order units, perhaps reflecting the abstract quality of the stimuli. Nevertheless, there were also striking differences between domains: For instance, during the temporal evolution of SL, responses to higher-order units continuously increased in the auditory domain, whereas responses in the visual domain developed faster and plateaued. This may reflect an artificial difference in the chunk sizes (pair vs. triplet) between the two modalities, or an innate difference in the learning curves between the auditory and visual learning pathways. In addition, the cortical areas involved in auditory and visual SL only partially overlapped (**fig. S9 and table S1)**. This was to be expected at the level of the sensory responses: responses encoding the sensory input and higher-order units clustered around the STG for the auditory SL involving syllables and around occipital cortex for the visual SL involving fractals. Perhaps more interestingly, areas engaged exclusively in higher-order unit representation were also partially separated. While areas such as IFG and ATL tracked higher-order units in both modalities, middle frontal and superior parietal cortices seemed to be differentially involved in auditory and visual SL, respectively. This result cannot be explained by the fact that different groups of subjects (and electrode coverage) contributed to the different tasks, as we confirmed this functional separation in six subjects who completed both auditory and visual SL tasks (fig. S9).

This suggests that while sequence operations performed across domains might build on similar representations (transitional probability, ordinal position and identity), the circuits performing these operations might be modality-specific to some extent, much like the nested tree structures involved in language, music, and mathematics that are each represented in distinct circuits. Our results speak for a more modularized representation of sequences for the encoding of local and simple aspects such as transitional probability represented in sensory areas, but a less modularized representation as complexity increases, as IFG and ATL encoded positional and identity information for both visual and auditory SL, in line with a domain-general role in SL. In turn, the hippocampus, at the top of the hierarchy, uniquely represents the identity of both visual and auditory sequences (*32*).

To our knowledge, the complementary representation of sequences across the cortex and the hippocampus, with a gradient of abstraction, has not been previously reported for SL. Our findings shed light on the elementary operations during SL and how cortex and hippocampus differentially support these processes. For instance, lower-level cortical coding based on transitional probabilities could facilitate initial segmentation, as uncertainty drives prediction errors and boundaries. Coding based on transitional probabilities, while a powerful cue to discover boundaries in the continuous stream, does not easily accommodate the integration and binding across elements. Higher-order cortical encoding based on ordinal position permits novel recombination of elements to create unique entities. In this case, so long ordinal position is respected, novel recombination of elements can be allowed. Finally, hippocampal integration and binding across stable combinations of units facilitates the attachment of meaning or identity. Thus, a great benefit of the observed complementary coding across the cortex and the hippocampus is that it may allow the further use of those information for different cognitive operations. These complementary roles of the cortex and the hippocampus were observed even during a brief exposure (∼10 mins). An interesting question for future research is to investigate the stability of these functions across longer exposure, and whether complementary coding persists or is replaced for a winner-take-all coding depending in the number of repetitions. Another question relates to how sleep consolidation affects these complementary representations.

An unexpected observation was the dissociation between ‘online’ and ‘offline’ behavioral measures of SL. Through frequency tagging we were able to localize with precision the areas involved in the acquisition of higher-order regularities in the structured stream — a response found in virtually all participants and that increased as a function of exposure. Furthermore, representational analysis demonstrated learning of transitional probability, ordinal position and unit identity after short exposure. Faster reaction times during the cover task showed that structured stream presentation facilitated learning behavior. Yet, the patients did not perform better than chance in the subsequent ‘offline’ behavioral recognition test. In neurotypical subjects, however, we observed facilitation in the reaction time in the structured stream and subsequent above-chance recognition performance in the offline behavioral task. In other words, cortical circuits for automatic SL appear intact in our patients, while episodic memory appear impaired. Indeed, episodic memory dysfunction is prevalent in temporal lobe epilepsy (*33*).

Relatedly, the fact that we observed SL in patients with epilepsy may challenge the importance of the medial temporal lobe (MTL) and hippocampus for SL. Previous fMRI studies in healthy subjects have demonstrated that visual SL leads to changes in representational similarity in the hippocampus (*9*); which we also observed in our population of patients. The critical role of the hippocampus has been corroborated by lesion studies in humans? of visual and auditory SL (*10, 25*). Computational models indicate a division of labor in the hippocampus between the monosynaptic pathway (connecting entorhinal cortex directly to CA1) which supports SL and the trisynaptic pathway (connecting entorhinal cortex to CA1 through dentate gyrus and CA3) which supports episodic memory (*34*). Given that epilepsy can lead to selective deficits in hippocampal circuits (*35*), the observed dissociation between online and offline behavioral measures of SL could reflect disproportionate damage to the trisynaptic pathway.

The dissociation between online and offline behavioral measures is also noteworthy given the large variability observed when SL is measured through explicit behavioral tasks (*36*). NFT may provide a more sensitive and robust measure of learning compared to explicit tasks, even in healthy populations. NFT can also be used to track learning in both the auditory and visual domains. This technique opens up exciting opportunities to characterize learning trajectories across clinical and healthy populations, across sensory modalities. Because NFT does require task demands, it is well-suited to tracking the acquisition of sequence knowledge across the lifespan from newborns to the elderly, and even in cognitively impaired patients. The combination of NFT with representational similarity analysis provides a powerful toolkit to reveal the how the brain engages in SL rapidly across multiple levels of organization in the human brain.

## Materials and Methods

### Stimulus materials and summary of experimental procedures

#### Auditory statistical learning task

Twelve consonant-vowel (CV) syllables were synthetically generated using MacTalk. Syllable lengths were equated and prosody was flattened using Praat (Boersma, Paul & Weenink, David, 2018). The individual syllables were concatenated in MATLAB. Two sequences were created: a structured and a random sequence. In the structured sequence transitional probabilities between syllables was manipulated such that 4 hidden words (3 syllables each) were embedded in the sequence (see **Fig. 1**), resulting in a continuous artificial language stream with an underlying syllable presentation rate of 4Hz, and word-rate of 1.33Hz. In the random sequence transitional probabilities across syllables were the same (e.g., p=1/11 syllables). Each sequence lasted approximately 2 minutes (540 syllable presentations) and was presented 5 times. To avoid potential cueing of the words at the start and end of the stream, the volume of the audio stream was ramped on and off, over the first and last 1.5s, respectively. Participants were not informed of the structure, and instead, to ensure task compliance, participants were asked to perform a cover task in which they indicated syllable repetitions that were randomly embedded in the auditory streams. Sixteen (*16*) syllable repetitions were randomly embedded into each presentation block of a sequence.

Once both streams (random and structured) had been played to the participants, they were then informed that one of the audio streams consisted of a hidden structured containing “words”. Subjects then performed a two-alternative forced-choice (2AFC) task where they had to select from two audio segments, presented one after the other, the one containing a “word”. One audio segment contained a “word” (e.g., “tupiro”), while the other was a lower probability “part-word” from the stream spanning word boundaries (e.g., “butupi”). Presentation order was counterbalanced across trials. Since exposure to the individual syllables is equated, a preference for the true words over the part-word is indicative of statistical learning. Two of the seventeen patients who participated in this experiment did not complete the 2AFC task, one for technical reasons and one because the participant was confused about the task.

#### Visual statistical learning task

The procedure for the visual SL task was identical to the auditory SL task, however, in this task, sequences were formed from 8 fractals (4-sets of two fractal pairs, duplets). Fractals were taken from the same set of images previously used in (*9*). The stimulus onset asynchorony between fractals was set to 375, whereby each fractal was presented for 233 ms with an interstimulus interval of 150 ms. In the structured sequence transitional probabilities between fractals were manipulated such that 4 hidden fractal-pairs (2 fractal each) were embedded in the sequence (see **Fig. 3**), resulting in a continuous stream of fractals with a presentation rate of 2.6 Hz and a fractal-pair rate of 1.3 Hz. In the random sequence, transitional probabilities remained fixed between all possible fractals (e.g., p=1/7). Each sequence lasted approximately 2 minutes (360 fractals presentations). As in the auditory learning task, participants were not informed of the structure, however, to ensure task compliance, participants were asked to perform a cover task in which they indicated, using the keyboard, when a fractal had been repeated. Sixteen (*16*) fractal repetitions were randomly embedded within each sequence block.

### Participants and recordings

#### Electrocorticography (ECoG)

ECoG recordings were obtained from a total of 23 patients (13 female, average age 35 yrs, range 16-59 yrs, 21 right-handed) with drug-resistant focal epilepsy undergoing clinically motivated invasive monitoring at the Comprehensive Epilepsy Center of the New York University Langone Medical Center. 11 subjects participated in the auditory statistical learning only, 6 in the visual statistical learning only, and 6 subjects participating both in the auditory and the visual statistical learning task. All subjects participating in the study provided oral and written informed consent prior to participation in the study, in accordance with the Institutional Review Board at the New York University Langone Medical Center. Patients were informed that participation in the study would not affect their clinical care and that they could withdraw from the study at any point without affecting medical treatment. Brain activity was recorded from a total of 3689 (average of 120+/-30 per subject) intracranially implanted subdural platinum-iridium electrodes embedded in silastic sheets (2.3 mm diameter contacts, Ad-Tech Medical Instrument). The decision to implant, electrode targeting, and the duration of invasive monitoring were determined solely on clinical grounds without reference to this or any other study. Macroelectrodes were arranged as grid arrays (8 × 8 contacts, 10 or 5 mm center-to-center spacing), linear strips (1 × 8/12 contacts), or depth electrodes (1 × 8/12 contacts), or some combination thereof. Subdural electrodes covered extensive portions of lateral and medial frontal, parietal, occipital, and temporal cortex of the left and/or right hemisphere (see **fig. S2** for electrode coverage across all subjects, and for the individual coverage of each subject). Recordings from grid, strip and depth electrode arrays were acquired using a NicoletOne C64 clinical amplifier (Natus Neurologics, Middleton, WI), bandpass filtered from 0.16-250 Hz and digitized at 512 Hz. Intracranial EEG signals were referenced to a two-contact subdural strip facing towards the skull near the craniotomy site. Data were subsequently downsampled to 250 Hz and a 60-Hz notch filter was applied to remove any line-noise artifacts. All electrodes were visually inspected, those with excessive noise artifacts were removed from subsequent analysis (185/3689 electrodes removed). In addition to analyzing the field potential (FP), high-gamma band (HGB) activity was extracted by applying an additional high-pass filter (fc=70 Hz) and the envelope of HGB activity was estimated by taking the square of the Hilbert transform of the filtered signal.

### Data analysis

#### ECoG Surface reconstruction and electrode localization

Pre-surgical and post-surgical T1-weighted MRIs were acquired for each patient, and the location of the electrode relative to the cortical surface was determined from co-registered MRIs following the procedure described in Yang and colleagues(*37*). Co-registered, skull-stripped T1 images were nonlinearly registered to an MNI-152 template and electrode locations were then extracted in Montreal Neurological Institute (MNI) space (projected to the surface) using the co-registered image. A three-dimensional reconstruction of each patient’s brain was computed using FreeSurfer (http://surfer.nmr.mgh.harvard.edu). In all figures, electrode locations are projected onto the left hemisphere of the MNI-152 template brain, unless otherwise noted.

### Behavioral data analysis

Performance during the online incidental task was assessed by calculating a d’ score for every participant across all 5 exposure trials. As the incidental task was embedded in the continuous stream of auditory or visual stimuli, detection of a syllable or image repetition was deemed accurate (“hit”) if the participant made a keyboard response within 250-1500ms of the occurrence of the repetition (Results are robust to the selection of the response window, as comparable results were obtained using response windows up to 750ms, 1000ms, and 3000ms). All other keyboard responses outside the valid response window were deemed false alarms. Significance of d’ scores were assessed via a one-sample t-test and comparison of d’ scores across conditions (structured vs. random) was assessed with a paired t-test. Analysis of reaction times between conditions was assessed using a Wilcoxon two-sided paired signed rank test between the average reaction times per condition and participant. Performance on the offline 2AFC explicit recognition test was assessed by determining the percent of correctly identified “words” or “fractal-pairs”, and subjected to a Wilcoxon signed rank test against chance performance (50%).

### Phase coherence analysis

For each experiment, signals from all five (*5*) blocks of a sequence (structured or random) were concatenated and then reshaped into 10-word segments (10 words x 90 trials x electrodes) and converted into the frequency domain via FFT (0.134 Hz resolution). Phase coherence was computed for each electrode, *R*^2^ = [∑^*N*^ *cos* Ø]^2^ + [∑^*N*^ *sin* Ø]^2^ over the 90 trials (*38*). Significance of the response at each frequency of interest (e.g., 1.33Hz & 4Hz) was determined by comparing the magnitude of the coherence response to 1000 phase-shuffled surrogate datasets and then subjected to FDR correction across all electrodes.

### Phase coherence latency analysis

The response latency was computed for all electrodes with a significant phase coherence at the higher-order rate (i.e., word rate or pair rate, 1.33 Hz) across all blocks. To identify a time-point when each electrode exhibited a significant response (e.g., time to first significant response), phase coherence was computed on smaller segments of data (10 words x 9 trials), using a sliding window of 10 words. A nonparametric cluster-based permutation statistic was used to test for a significant change in coherence from baseline (e.g. random condition) across time (e.g. exceedance mass test on the sum of t-values in a cluster compared to 1000 permutations of the condition labels, (*39*)).

### Representational similarity analysis

To assess the similarity of neural responses to each token (syllables, fractals), a multivariate spatial pattern analysis was performed. First, all individual trials were ‘whitened’ by the noise covariance matrix computed across all tokens and trials(*40*) and the average response across all tokens (e.g., non-specific response) was subtracted from each trial. Next, the data were vectorized across all significant electrodes (e.g., samples x significant electrodes) within a cluster (e.g., word+syll or word-only) and the dissimilarity of the spatial patterns was computed between each pair of tokens. Dissimilarity was assessed using the correlation distance, 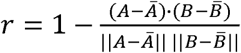, where A and B represent the mean spatial vectors for two of the tokens being compared (e.g., syllables ‘tu’ vs. ‘pi’), and using 5-fold cross-validation. The resultant representational dissimilarity matrix (RDM) was subjected to a principal component analysis and the first two dimensions were plotted against each other to produce a 2-dimensional mapping of dissimilarity scores across all pairs (see **Fig. 2**).

### Within vs. between category similarity analysis

To quantify the degree to which responses are able to capture features of the learned streams i.e., transitional probability, ordinal position and/or identity, we calculated the difference in similarity (Pearson correlation) between items in the same category and items belonging to the other category in question. This similarity estimate was calculated by randomly sampling the significant electrodes by type (e.g., FP, word-only) and computing a similarity matrix between all tokens (e.g., syllables or fractals), for each resampling. This resampling procedure was repeated 200 times (with replacement), and the average fisher-transformed correlations of all elements within a category was compared against all items that spanned the opposing category using a Wilcoxon’s rank sum test (two-sided).

## Acknowledgements

We thank P. Minhas, B. Mahmood, M. Hofstadter for help in intracranial data collection, H. Wang for electrode reconstruction, Sasha Devore, Bijan Pesaran, David Poeppel and Caspar M. Schwiedrzik for helpful comments on an earlier version of the manuscript. This work was supported by a grant from the German-Israeli Foundation for Scientific Research and Development (GIF I-99-105.4-2016) to L.M., and NIH R01 MH069456 and the Canadian Institute for Advanced Research to N.B.T-B. The funders had no role in study design, data collection and analysis, decision to publish, or preparation of the manuscript. The authors declare no competing interests.

## Author Contributions

Conceived and designed the experiments: LM; collected the data: SH and LM.; attending neurosurgeons: WD; supervised recordings and IRB approval process: AF, DF, OD, WD, PD and AL; analyzed the data: SH, NTB and LM; consulted on experimental design and interpretation: NTB; wrote the manuscript: SH, NTB and LM; revised the manuscript: all authors.

## Competing Interests

The authors declare no competing interests.

## Data and materials availability

The data sets generated and code used to generate the main findings of the current study are available from the corresponding author upon reasonable request.

## Notes

### Competing Interest Statement

The authors have declared no competing interest.

### Summary of Updates

New results are presented from hippocampus; The title has been updated to reflect the broader scope of the article and results. Supplemental files updated

