## Supplemental Data for "Learning hierarchical sequence representations across human cortex and hippocampus"

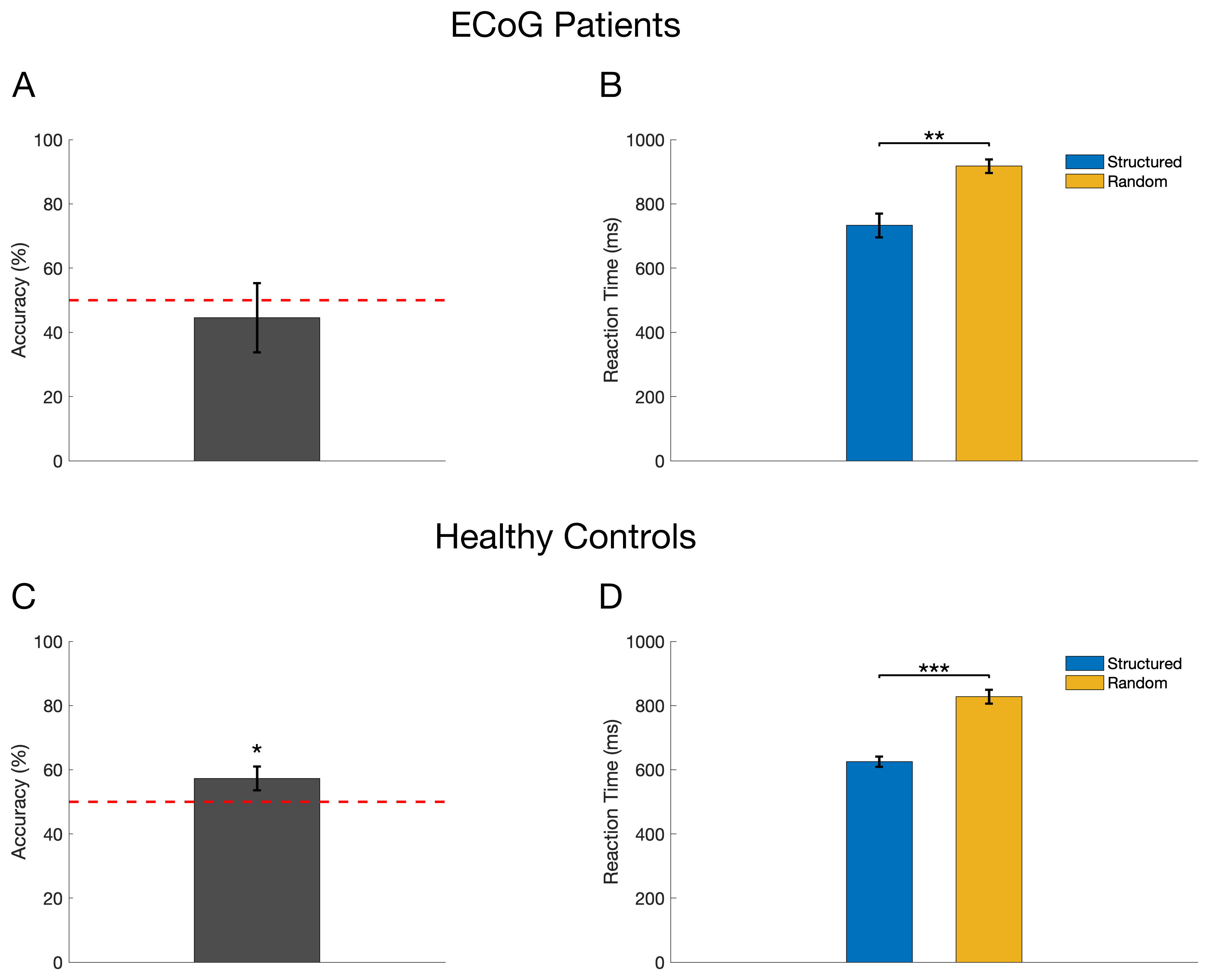

**Supplementary Figure S1.** Behavioral results in the auditory SL task for surgical ECoG patients (**Top row**) and healthy controls (**Bottom row**). **(A, C)** Accuracy (percent correct) on the offline, explicit recognition task. (**B, D**) Reaction times associated with the online, 1-back cover task (e.g. syllable repetition task). While ECoG patients were not able to explicitly recall the real-words vs. part-words (t(16)=-1.9, p=0.07), they did exhibit faster reaction times to syllable repetitions in the structured vs. random streams (p<0.01), comparable to those exhibited by healthy controls that did show explicit recognition (t(16)=2.1, p=0.04).

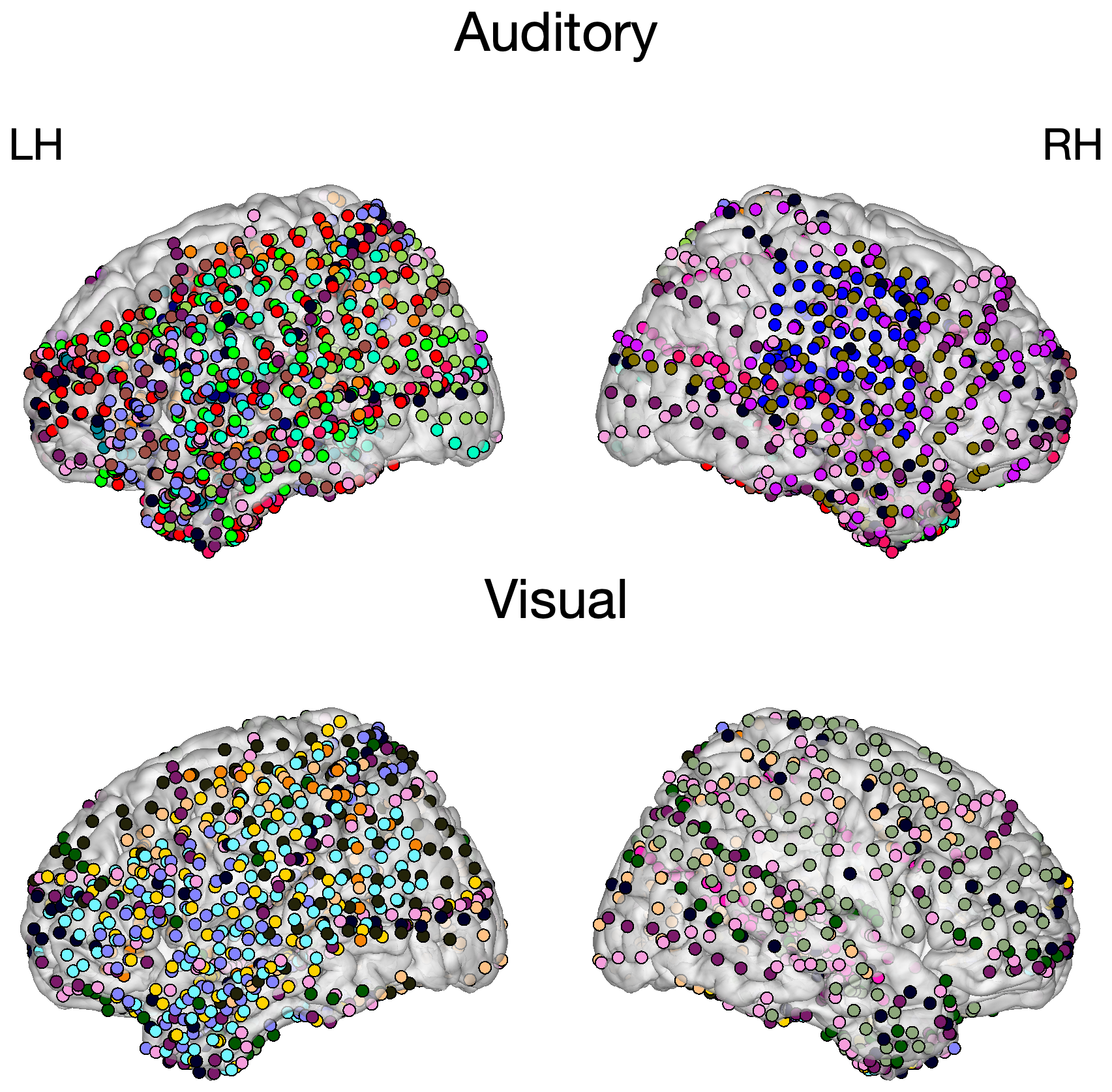

**Supplementary Figure S2.** ECoG electrode coverage for participants in the auditory SL task (Top) and visual SL task (Bottom). Each participant’s electrode coverage is shown in a different color on the left hemisphere (LH) and right hemisphere (RH).

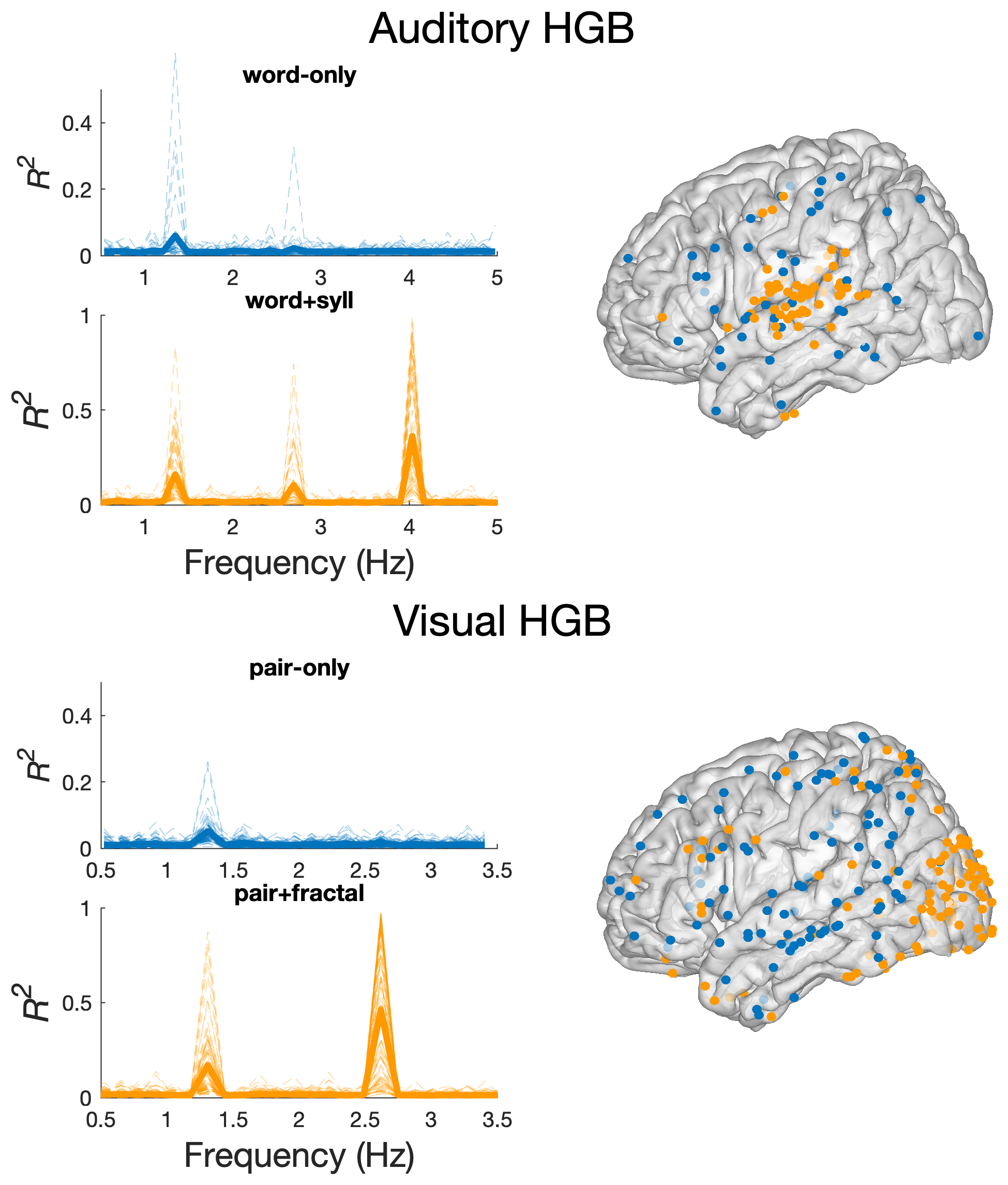

**Supplementary Figure S3.** High-gamma band **(**HGB) phase coherence analysis of the auditory (**Top**) and visual (**Bottom**) SL tasks. Electrodes showing a significant response in the HGB envelope at the word-only / pair-only rate are shown in blue. Electrodes showing a significant response at the word+syll / pair+fractal rate are shown in orange (*p*<0.05, FDR corrected).

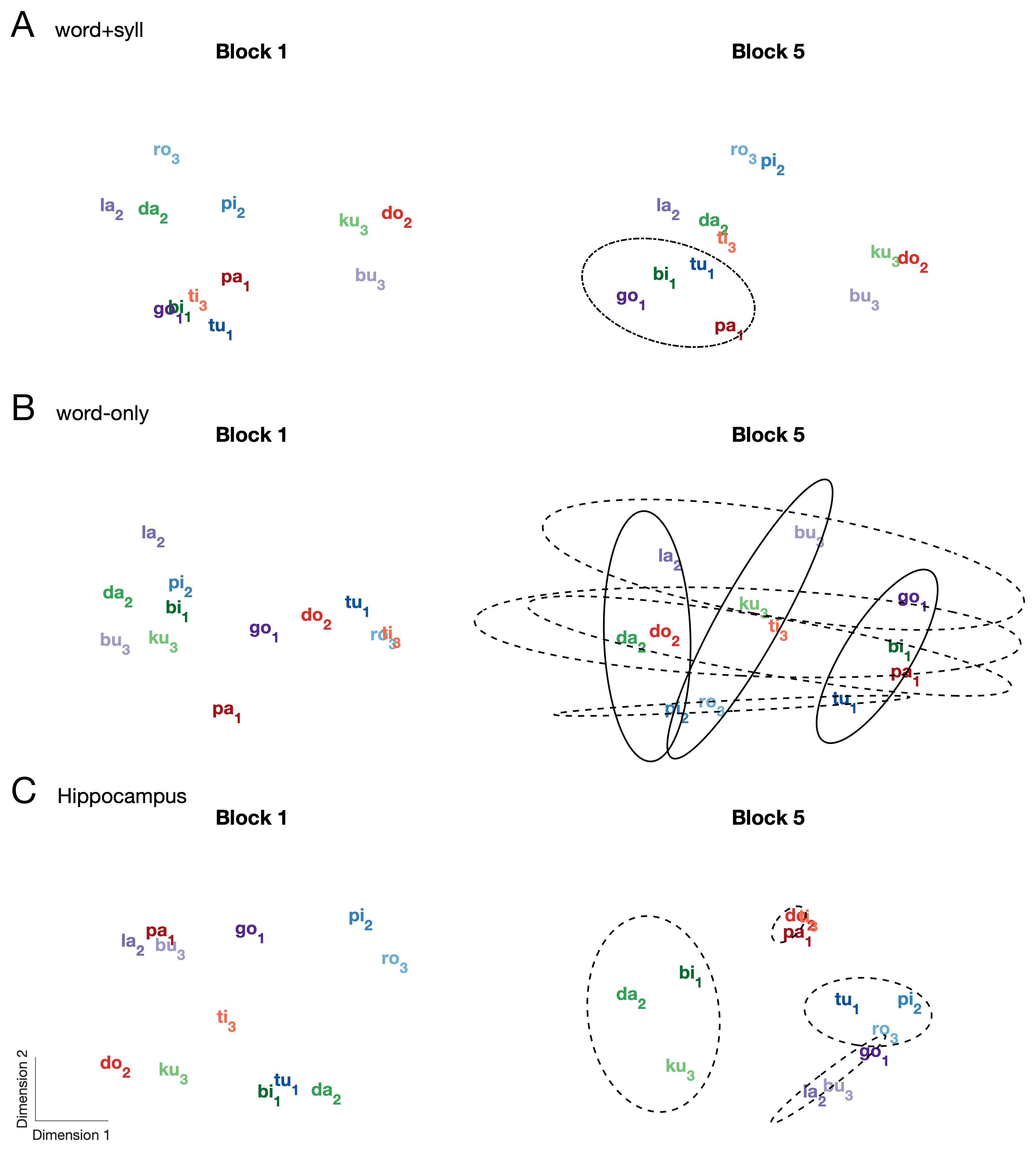

**Supplementary Figure S4.** Representational similarity analysis (RSA) across 1^st^ vs. 5^th^ blocks during the auditory SL task for different electrode types. Multidimensional scaling (MDS) shows that different coding schemes emerge by the 5^th^ block and are not present in the initial block.

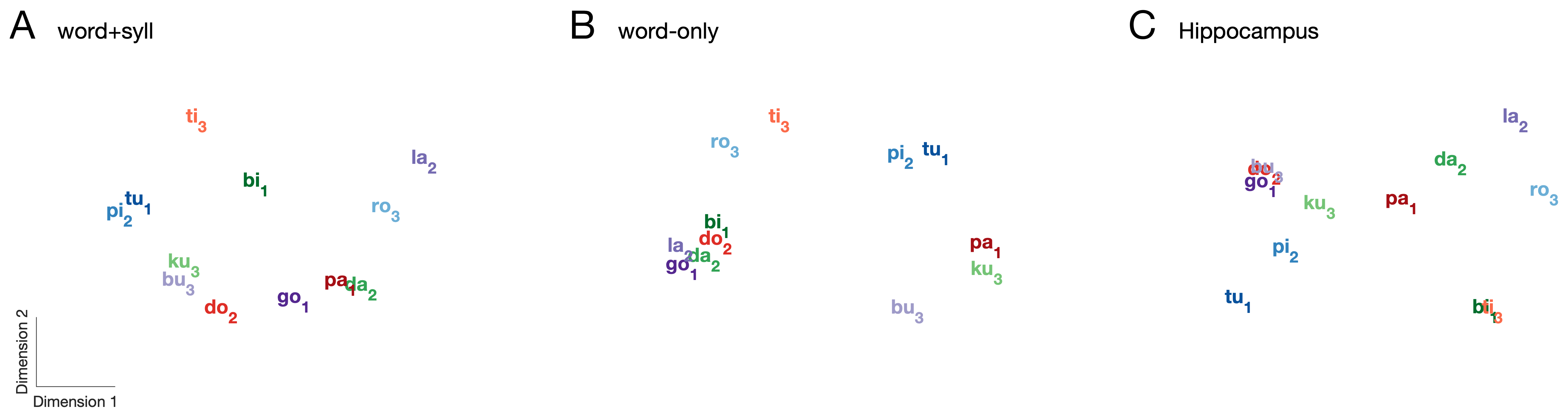

**Supplementary Figure S5.** Representation similarity analysis (RSA) of syllables during random sequence in the auditory SL task. Multidimensional scaling (MDS) of distances between responses to individual syllables for electrodes showing significant HGB coherence in (A) word+syll and (B) word-only electrodes, and (C) Hippocampal electrodes. MDS of distances between syllable responses did not show any clustering based on transitional probability, ordinal position, or word identity for any electrode type.

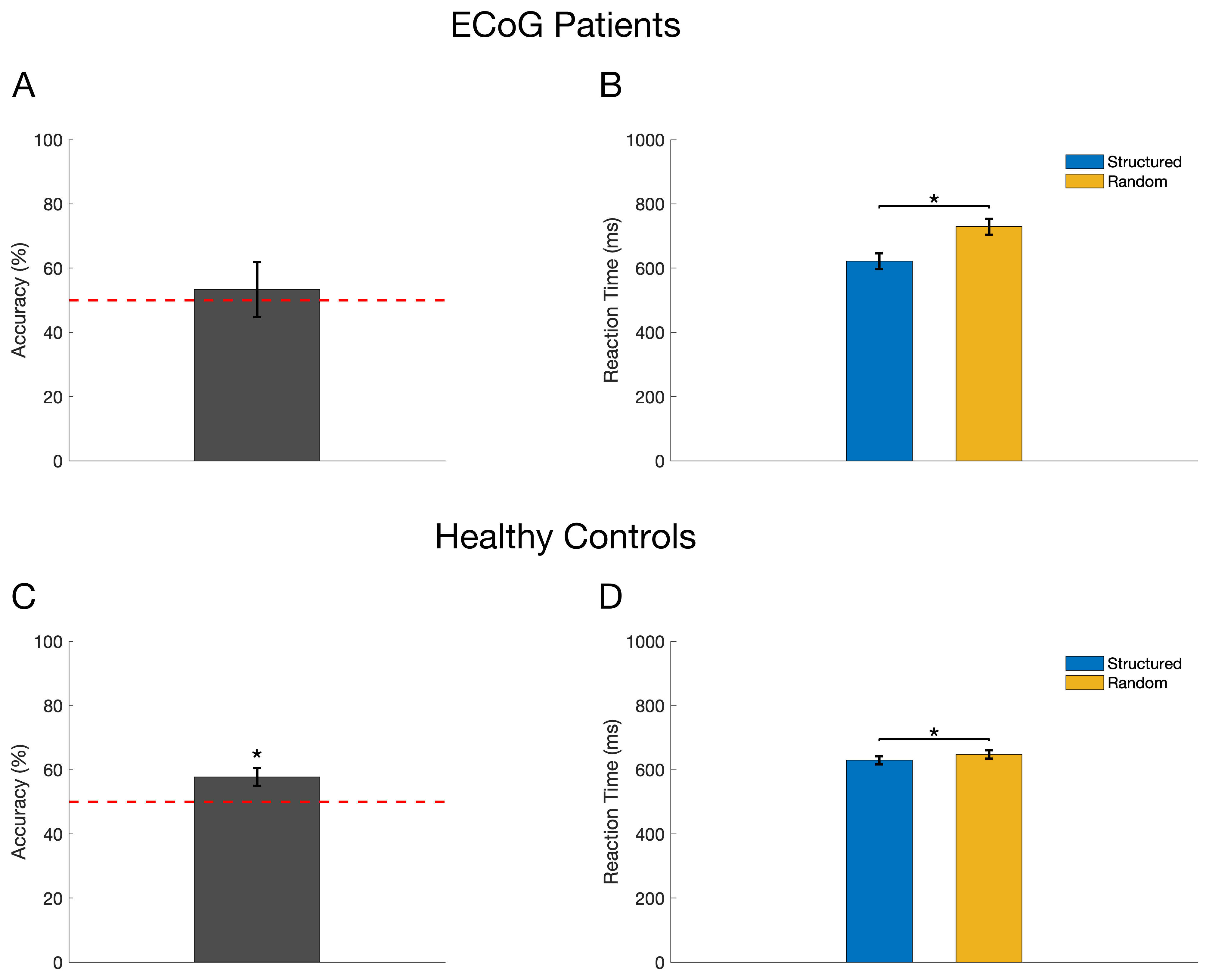

**Supplementary Figure S6.** Behavioral results in the visual SL task for surgical ECoG patients (**Top row**) and healthy controls (**Bottom row**). **(A, C)** Accuracy (percent correct) on the offline, explicit recognition task. (**B, D**) Reaction times associated with the online, 1-back cover task (e.g. fractal repetition task). As in the auditory SL task, ECoG patients were not able to explicitly recall the real-pairs vs. part-pairs (t(12)=1.4, p=0.18), they did exhibit faster reaction times to fractal repetitions in the structured vs. random streams (p<0.05), comparable to those exhibited by healthy controls that did show explicit recognition (t(13)=2.89, p=0.01).

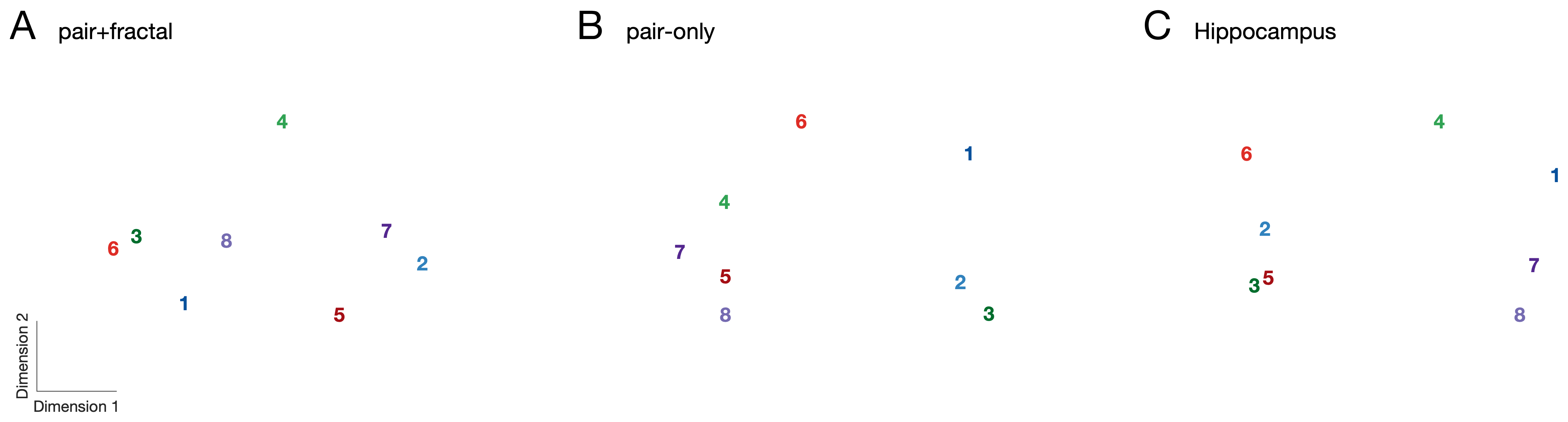

**Supplementary Figure S7.** Representation similarity analysis (RSA) of fractals during random sequence in the visual SL task. Multidimensional scaling (MDS) of distances between responses to individual fractals for electrodes showing significant HGB coherence in (A) pair+fractal, (B) pair-only and (C) Hippocampal electrodes. Pairs are color-coded, odd numbers refer to the first position, even to the second. MDS of distances between fractal responses did not show any clustering based on transitional probability, ordinal position, or pair identity for any electrode type.

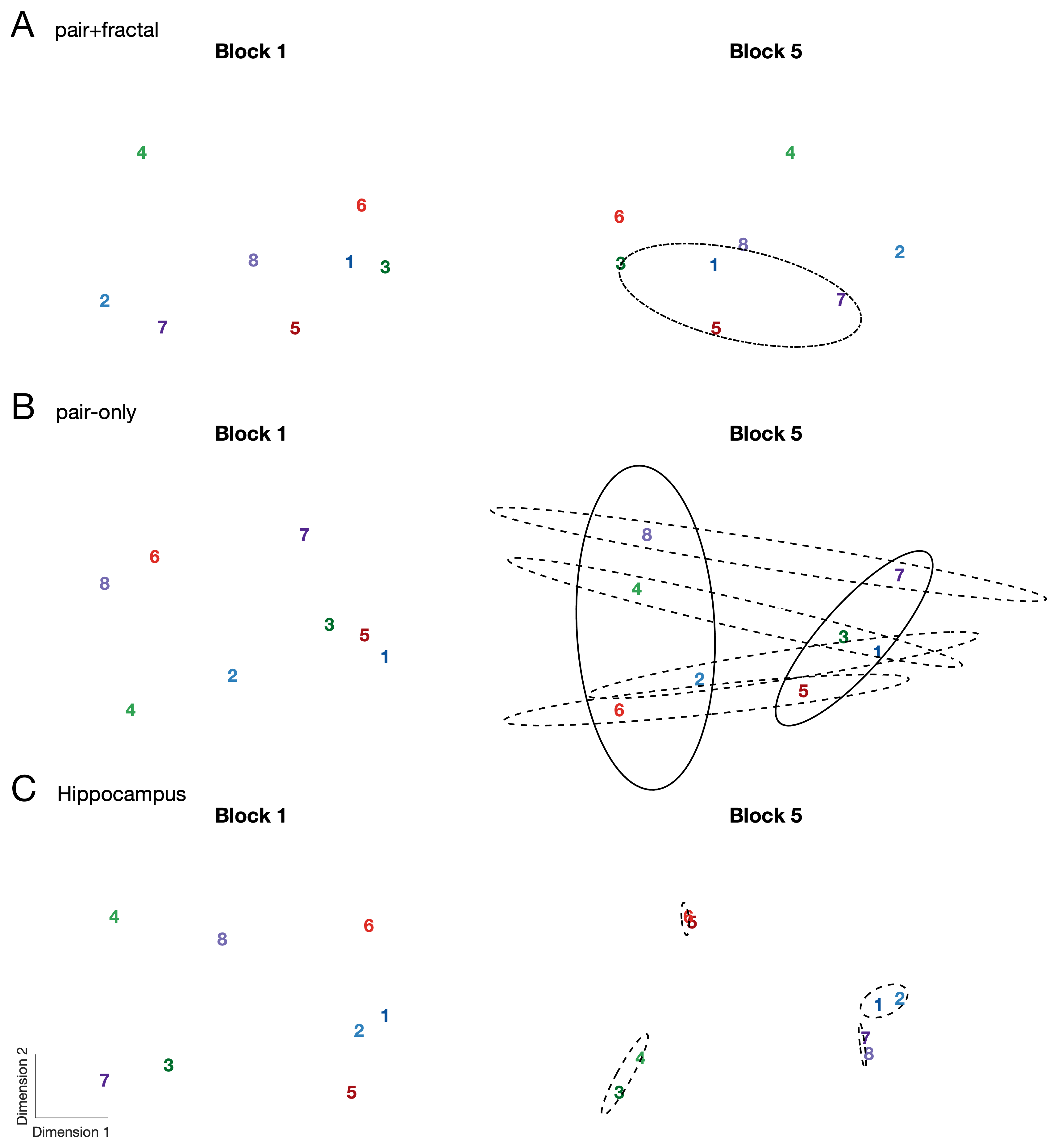

**Supplementary Figure S8.** Representational similarity analysis (RSA) across 1^st^ vs. 5^th^ blocks during the visual SL task for different electrode types. Multidimensional scaling (MDS) shows that different coding schemes emerge by the 5^th^ block and are not present in the initial block.

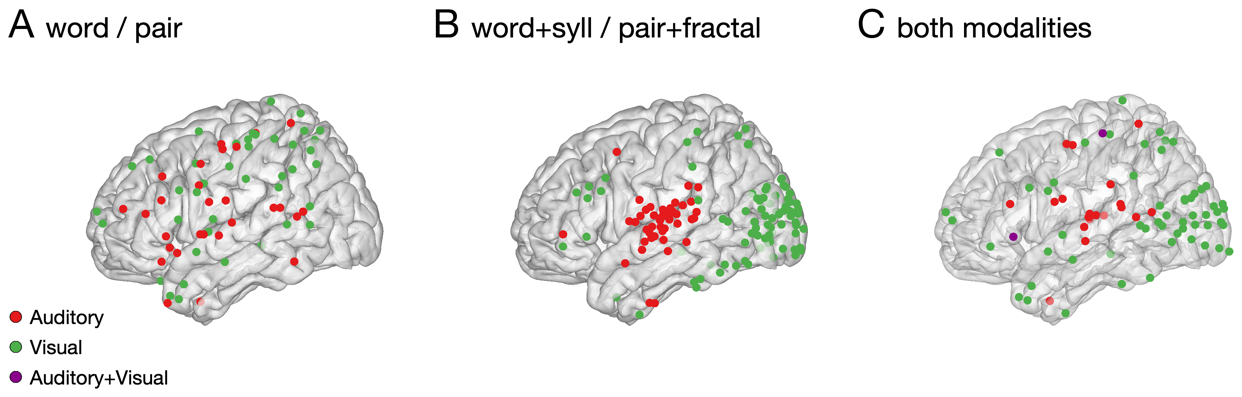

**Supplemental Figure S9. Task overlap in NFT.** (**A-B**) Electrode localization, projected onto the left hemisphere, of higher-order structure in the HGB across auditory (red) and visual (green) SL. (**C**) Electrode overlap across modalities for six subjects who participated in both tasks. Electrodes exhibiting a significant response in the auditory (Red) and visual (Green) SL tasks. Purple electrodes depict electrodes that were found to have a significant response in both tasks. In all, 2 electrodes were found to have significant responses in both modalities across the six subjects. Electrodes are projected onto the left hemisphere for display purposes.

|  | **Word-only / Pair-only** | | **Word+Syll / Pair+Fractal** | |
| --- | --- | --- | --- | --- |
| **ROI** | **Auditory** | **Visual** | **Auditory** | **Visual** |
| bankssts | 1 | 0 | 1 | 4 |
| caudalmiddlefrontal | 1 | 6 | 3 | 12 |
| cuneus | 0 | 1 | 0 | 6 |
| entorhinal | 0 | 1 | 2 | 3 |
| fusiform | 1 | 4 | 2 | 17 |
| inferiorparietal | 1 | 7 | 3 | 31 |
| inferiortemporal | 0 | 6 | 2 | 6 |
| isthmuscingulate | 1 | 1 | 0 | 2 |
| lateraloccipital | 1 | 3 | 2 | 35 |
| lateralorbitofrontal | 0 | 1 | 0 | 2 |
| lingual | 0 | 0 | 0 | 8 |
| medialorbitofrontal | 0 | 0 | 0 | 1 |
| middletemporal | 2 | 2 | 1 | 5 |
| parahippocampal | 0 | 0 | 0 | 1 |
| paracentral | 0 | 1 | 0 | 1 |
| parsopercularis | 1 | 3 | 7 | 5 |
| parstriangularis | 3 | 5 | 2 | 1 |
| pericalcarine | 0 | 0 | 0 | 3 |
| postcentral | 0 | 13 | 17 | 14 |
| posteriorcingulate | 0 | 0 | 0 | 2 |
| precentral | 2 | 6 | 21 | 17 |
| precuneus | 0 | 1 | 0 | 7 |
| rostralmiddlefrontal | 3 | 9 | 1 | 13 |
| superiorfrontal | 0 | 0 | 0 | 7 |
| superiorparietal | 1 | 7 | 2 | 41 |
| superiortemporal | 0 | 1 | 14 | 10 |
| supramarginal | 0 | 15 | 13 | 32 |
| temporalpole | 2 | 3 | 2 | 0 |
| transversetemporal | 1 | 0 | 5 | 2 |
| cMTG | 0 | 3 | 1 | 8 |
| mMTG | 0 | 4 | 2 | 2 |
| rMTG | 2 | 3 | 2 | 2 |
| cSTG | 1 | 1 | 18 | 11 |
| mSTG | 1 | 4 | 10 | 2 |
| rSTG | 1 | 2 | 2 | 0 |
| amyg | 1 | 4 | 1 | 22 |
| insula | 0 | 0 | 14 | 20 |
| uncertain localization | 0 | 6 | 15 | 77 |

**Supplemental Table S1.** Number of significant FP electrodes exhibiting phase coherence in different anatomical regions broken down by response type (e.g., word-only, word+syll).
